# TF2TG: an online resource mining the potential gene targets of transcription factors in *Drosophila*

**DOI:** 10.1101/2025.02.13.638157

**Authors:** Yanhui Hu, Jonathan Rodiger, Yifang Liu, Chenxi Gao, Ying Liu, Mujeeb Qadiri, Austin Veal, Martha L. Bulyk, Norbert Perrimon

## Abstract

Sequence-specific transcription factors (TFs) are key regulators of many biological processes, controlling the expression of their target genes through binding to the cis-regulatory regions such as promoters and enhancers. Each TF has unique DNA binding site motifs, and large-scale experiments have been conducted to characterize TF–DNA binding preferences. However, no comprehensive resource currently integrates these datasets for *Drosophila*. To address this need, we developed TF2TG (“transcription factor to target gene”), a comprehensive resource that combines both *in vitro* and *in vivo* datasets to link transcription factors (TFs) to their target genes based on TF–DNA binding preferences along with the protein-protein interaction data, tissue-specific transcriptomic data, and chromatin accessibility data. Although the genome offers numerous potential binding sites for each TF, only a subset is actually bound *in vivo*, and of these, only a fraction is functionally relevant. For instance, some TFs bind to their specific sites due to synergistic interactions with other factors nearby. This integration provides users with a comprehensive list of potential candidates as well as aids users in ranking candidate genes and determining condition-specific TF binding for studying transcriptional regulation in *Drosophila*.

## Introduction

Transcriptional regulation is important for many biological processes including cell growth, differentiation, and responding to environmental changes. Transcription factors (TFs) regulate the expression of specific genes by binding to genomic sequences in regions such as promoters or enhancers. Binding of TFs to DNAs can recruit other proteins and complexes that either aid or obstruct the transcriptional process, which are crucial in controlling whether the downstream genes are turned on or off. Mis-regulation in TFs is frequently observed in diseases, including cancer and genetic disorders.

The methods to characterize TF–DNA binding preferences can be broadly categorized into *in vivo* and *in vitro* approaches. TF binding motifs are typically short, conserved DNA sequences that are recognized by particular TFs and are primarily found in noncoding intergenic or intronic DNA, with a few exceptions (Stergachis *et al*. 2013). TF binding motifs have predominantly been identified through *in vitro* methods. Early studies used biochemical methods such as DNase I foot-printing (Bergman *et al*. 2005), while SELEX (Systematic Evolution of Ligands by EXponential enrichment) (Roulet *et al*. 2002), PBMs (Protein Binding Microarrays) (Mukherjee *et al*. 2004) and B1H (bacterial 1 hybrid) assays (Zhu *et al*. 2011) have been used subsequently for more systematic evaluation of numerous potential target sequences for a single TF with higher resolution. Several existing databases can be mined for TF DNA binding motifs in *Drosophila* such as the commercial database of Transfac (Wingender 2008), the JASPAR database (Rauluseviciute *et al*. 2024), the UniPROBE database (Hume *et al*. 2015) and FlyFactorSurvey (Zhu *et al*. 2011), as well as an integrated resource OnTheFly (Shazman *et al*. 2014).

Although the genome contains numerous potential binding sites for each TF, only some of them are bound *in vivo* and the binding sites might vary under different biological contexts. In recent years, a number of efforts and resources have built technologies and generated large amounts of data to identify TF binding *in vivo*. The most predominant *in vivo* technique to detect TF–DNA binding is chromatin immunoprecipitation combined with high-throughput sequencing (ChIP-seq). In this method, genomic regions associated with a specific TF are isolated using immunoprecipitation, and the bound sequences are then identified through high-throughput sequencing. For *Drosophila*, much of the work defining TF-chromatin interactions has been accomplished by the modENCODE/modERN (model organism Encyclopedia of Regulatory Networks) consortium (Kudron *et al*. 2018; Kudron *et al*. 2024). Specifically, GFP-tagged strains covering 658 TFs were made and grown to the desired stage for whole animal ChIP-seq. Data of ChIP-seq binding profiles for 604 TFs (more than 90% of the fly TFs) with a total of 674 experiments have been successfully generated using a highly specific anti-GFP antibody, resulting in more than 3.6M peaks (78K meta-peaks) in the fly genome (Kudron *et al*. 2018; Kudron *et al*. 2024). The modERN consortium recently re-processed all ChIP-seq datasets using a newer version of the analysis pipeline and provides an online resource for users to retrieve peak analysis results for downstream analysis and visualization. The REDfly resource is another important resource providing curated cis-regulatory modules (CRMs) as well as genomic TF binding sites for *Drosophila* from the literature (Keranen *et al*. 2022).

Here, we developed TF2TG to integrate multiple lines of evidence on transcriptional regulation including the potential genomic TF binding sites based on the TF binding motifs identified *in vitro*, the peak signals from *in vivo* data obtained based on ChIP-seq technology by the modEncode/modERN consortium, manual curation of CRMs from REDFly as well as the high-throughput chromosome conformation capture data (Hi-C) (Dekker *et al*. 2017). TF2TG allows researchers to inquire: What TFs might bind to the promoter region of a given gene? What TF peaks are observed in the promoter region of a given gene and in a certain context, based on ChIP-seq or ChIP-chip data? What are the potential long-distance enhancers that might regulate expression of the gene? Currently, there is no public resource that can be easily mined to answer these types of questions. TF2TG makes it possible to use high-throughput and other TF binding site-related and transcriptomics data sets to predict what TFs might regulate a given gene of interest. By further integrating protein-protein interaction data as well as tissue/cell type specific expression data, TF2TG can help to identify context specific TFs that are functionally related. As increasing expression profiles and phenotypic data become available to identify genes with key roles in various biological processes, TF2TG provides a valuable resource for investigating the underlying transcriptional regulatory networks.

## MATERIAL AND METHODS

### 1. Motif scan

DNA binding site position weight matrices (PWMs) for *Drosophila* TFs were obtained from FlyFactorSurvey data (https://mccb.umassmed.edu/ffs/) (Zhu *et al*. 2011) as well as the curated/filtered PWMs from publication (Shokri *et al*. 2019). We used MOODS (Korhonen *et al*. 2009), a suite of advanced matrix matching algorithms (https://github.com/jhkorhonen/MOODS), for scanning Drosophila genomic sequences with PWMs. Then the target genes were identified if the motif site is located upstream of the transcription start site (TSS) or the intron regions of any genes with 4 different search windows (1 kb,2kb, 5kb and 10 kb).

### 2. ChIP-seq datasets

The raw data files of 48 datasets curated from publication (Shokri *et al*. 2019) were obtained from GEO and were processed using a local pipeline to align reads with the reference genome using Bowtie2. After filtering using Samtools and Sambamba, peak calling was performed using MACS2. In addition, the processed peak information of 660 datasets from modERN (https://epic.gs.washington.edu/modERNresource/) were directly used and integrated. Then the target genes were identified if the middle point of the peak is located upstream of the transcription start site (TSS) or the intron regions of any genes with 4 different search windows (1 kb,2kb, 5kb and 10 kb).

### 3. Hi-C data

The data files of in situ Hi-C on *Drosophila* S2 cells were obtained from the 4DN data portal (https://data.4dnucleome.org/resources/data-analysis/hi_c-processing-pipeline) (Dekker *et al*. 2017) and processed using Juicer (Durand *et al*. 2016). The raw interaction matrix was normalized by HiCNorm to correct the GC content in different DNA regions, effective length, and differences in comparison rate. After obtaining a genome-wide interaction map, downstream detection of 3D structure included compartment A/B structure detection based on principal component analysis, TAD structure detection based on Markov model, and loop structure detection based on peak measurement using the Juicer tool HiCCUPS.

### 4. Other datasets and annotation

Protein and protein interaction data as well as genetic interaction data were obtained from MIST database (Hu *et al*. 2018). Tissue-specific RNA-seq data were obtained from the DGET database (Hu *et al*. 2017). CRMs annotated by REDfly (Keranen *et al*. 2022) were obtained from FlyBase. ATAC-seq datasets were obtained from GEO (Gene Expression Omnibus Database) (Clough and barrett 2016) (sup table 1). The transcriptomic data of Yorkie (Yki) gut from day 8 adult flies was also obtained from GEO (GSE113728).

## Results

### 1. Data integration

The field of regulatory genomics is generating a vast amount of data, including sequencing data related to TF binding (e.g. ChIP-seq data from ENCODE and modEncode), Hi-C data related to long-distance enhancers (e.g. 4DN Hi-C data), protein-protein interaction data among TFs (e.g. as integrated at MIST), and transcriptional profile data (e.g. modEncode). Building a resource that integrates these data sets, together with computational prediction of TF binding, will help researchers use different types of information to gain a more complete understanding of gene regulation based on transcriptional networks and prioritize genes based on integrated information (Figure 1).

**Figure 1:**
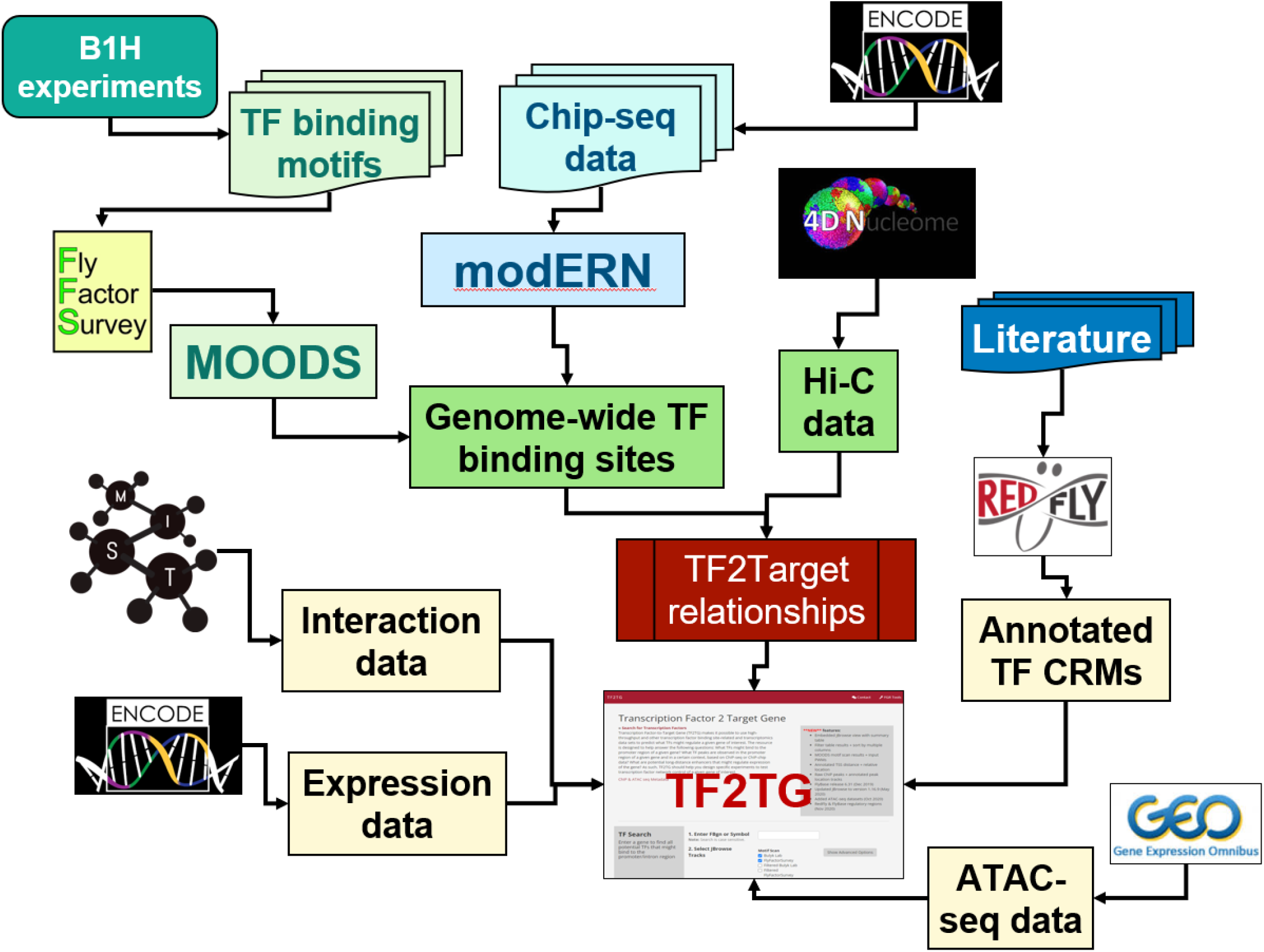
Design schema of TF2TG tool. TF2TG was developed to integrate multiple lines of evidence on transcriptional regulation including the potential genomic TF binding sites based on TF binding motifs identified *in vitro*, the peak signals from *in vivo* data obtained based on ChIP-seq data from modEncode/modERN consortium, manual curation of CRMs from REDFly, ATACseq datasets from GEO, as well as the high-throughput chromosome conformation capture data (Hi-C) along with tissue specific transcriptomic data and protein-protein interaction data.

TFs can display binding specificity for multiple, distinct nucleotide sequence motifs typically of 6-12 bases in length. Motifs have been identified largely using *in vitro* methods that can systematically assess many possible target sequences for a single TF. Among the publicly available database resources focusing on *Drosophila* TF binding motifs, FlyFactorSurvey houses more than 400 TF DNA binding motifs for 327 TFs, primarily obtained from a B1H assay (bacteria 1 hybrid assay) supplemented with literature annotation and more importantly it also provides the ftp file of all the TF binding motifs in various file formats. Although FlyFactorSurvey, as well as other similar resources, provides the ftp file of all the TF binding motifs in various file formats and a web-based interface for users to mine the TF binding motif, it is not trivial to convert the information of binding preference into unambiguous sequence information since the binding sites sometimes are quite degenerate. To help bench researchers mine the information, we downloaded the TF motifs in PWM format from FlyFactorSurvey and ran the motif finding algorithm MOODS (Korhonen *et al*. 2009) to align the motifs on the *Drosophila* genome to identify all the potential sites for these 327 TFs (TF A, Figure2). While more than 50% of TFs lack binding motif PWMs, and not all of the TF binding site motif occurrences on chromosomes are used *in vivo*, it is important to include the datasets obtained from *in vivo* experiments as well (TF B and C, Figure2).

**Figure 2:**
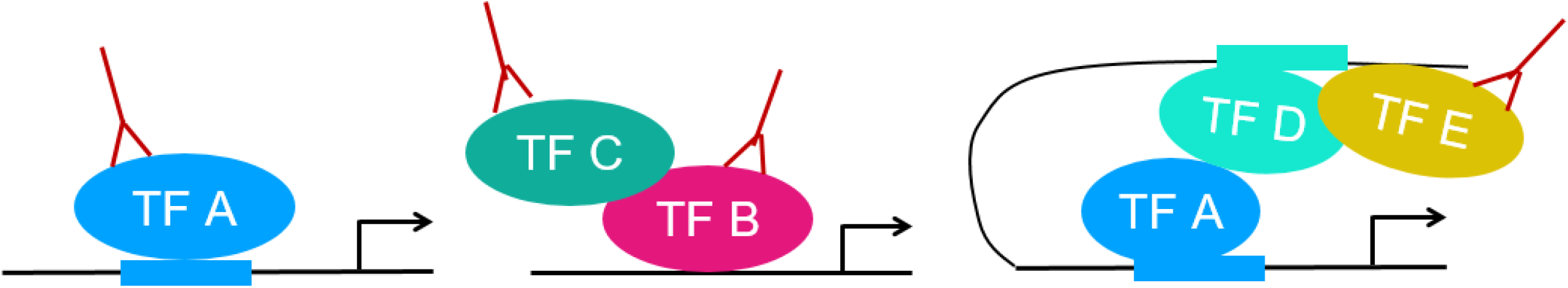
Scenarios of TF binding sites. Binding motif information is available for TF A but not for TF B. TF C is a cofactor for TF B binding, while TF D and TF E bind at a distance enhancer region identified by Hi-C data.

The modENCODE/modERN consortium generated hundreds of GFP-tagged strains covering more than 90% of TFs for *Drosophila*. The expression patterns of the TFs were characterized and the whole animal ChIP-seq analysis was performed at the stage when the corresponding TF is well expressed. Specifically, the majority of TFs (525 experiments) were done using embryo samples, while 126 larva/pupa samples and 22 adult samples were used for the remaining TFs. Overall, ChIP-seq was successfully done for 604 fly TFs and 56 other DNA-associated proteins. The datasets were generated and analyzed over a decade and recently the modERN consortium reanalyzed all the datasets through an improved peak calling pipeline using more recent releases of the relevant tools (SPP and ENCODE-DCC pipeline). In these uniformly processed datasets, more than 3.6 million peaks were identified in the *Drosophila* genome. After the clusters of peaks were consolidated using MACS2, about 78K megapeaks were identified for *Drosophila*. Further, the modERN consortium made the information of optimal peaks identified from the reanalysis effort available for download at the modERN website (Kudron *et al*. 2024).

We developed a pipeline to map both the motif scan results and the peaks from ChIP-seq relative to annotated genes on the chromosome and inferred the TF-target gene relationships if the TF binding sites and/or ChIP-seq peak are present in the promoter of the transcriptional start site (TSS) and/or the intron regions of target gene. In addition, TF2TG also takes into consideration the manual curation of CRMs from REDfly as well as the topologically associating domains (TADs) in proximity to a given promoter region in 3-D space (Figure 1), which can help to identify TFs that bind to a long-distance enhancer (TF D and E, Figure2).

### 2. UI features

We implemented TF2TG, an online resource for users to mine the integrated information from motif scans, peak calling from ChIP-seq data and in situ Hi-C data. There are 3 options at the user-interface. First, user can input any gene to look for the potential TFs that might regulate that gene. All the TFs with binding motif(s) or ChIP-seq peak(s) identified within the search space that the user specifies, such as one thousand base pairs upstream and downstream from transcription starting site (TSS), are summarized in a result table as well as visualized using JBrowser (sup figure 1). In this summary table, protein-protein interaction and genetic-interaction data among all the potential TFs within the results table are also provided. In addition, users have the option to build a node-edge graphical network of all the potential TFs identified based on either all protein-protein interaction data or high-quality binary interaction data from MIST (Hu *et al*. 2018; Tang *et al*. 2023). Users can also choose a tissue of interest and use the coloring of gene nodes to show the expression level of relevant TFs on the network (sup figure 1). The second search option on the TF2TG landing page is for users to input a TF and identify all the potential target genes. All the potential target genes with the motif binding and/or peaks sites bound by the input TF are summarized in a results table. For both search options, tissue-specific RNA-seq datasets are provided in the result page and a user can select a tissue of interest to prioritize the candidate genes based on their expression level. The options of search space at TF2TG is 1k, 2k, 5k or 10k upstream and downstream of TSS masking exon regions. Users can further refine the results using more stringent criteria since the location information as well as the distance to TSS are provided in the results table. We also integrated data on open chromatin regions from more than 500 ATAC-seq datasets submitted to GEO from 30 publications (sup table 1) making it possible for user to refine the result by selecting the relevant ATAC-seq dataset (sup figure 1). Users also have the option in TF2TG to search based on curated TF motifs and/or filtered TF motifs as well as a subset of ChIP-seq datasets (Shokri *et al*. 2019). The third option on the search page is for a user to input a genomic location and retrieve all the associated TADs identified by in situ Hi-C data. In addition, the motifs and peaks identified in the associated TADs are displayed on the results page. For all the three search options, the detailed table of motif scan, e.g. the motif sequence, motif location, and motif score are provided, as well as the detailed table of peak information from ChIP-seq data such as the location of the peaks and the sample information.

### 3. Use case

As an example, we used TF2TG to identify putative TFs that regulate factors secreted by tumors. Understanding the regulatory mechanisms of tumor-secreted factors is critical for elucidating how tumors communicate with host organs and disrupt their physiological functions. In a previously established fly model of organ wasting/cancer cachexia, it was demonstrated that adult gut tumors, induced by overexpression of an activated form of Yorkie (Yki)(referred to as Yki tumors), secrete several factors, including ImpL2 (Ecdysone-inducible gene L2), PDGF- and Pvf1 (VEGF-related factor 1), and upd3 (unpaired 3), which leads to many cachexia-like phenotypes such as muscle wasting (Kwon *et al*. 2015; Song *et al*. 2019; Ding *et al*. 2021). However, the TFs regulating the expression of these secreted factors remain uncharacterized. Thus, we searched the TF2TG database for TFs that could potentially regulate each ligand with a 5k search window (5k base pair upstream of TSS and intron regions). Combining the TFs for each secreted factor resulted in a list of 640 TFs (Figure 3A, sup table 2), with 361 TFs identified by motif scan, 621 from ChIP-seq data. 171 TFs were identified both by motif scan and ChIP-seq data (Figure 3B), which is expected since not all the TFs have binding motif data available. We observed significant overlaps among the TFs for the secreted factors, with 197 TFs (31%) shared by all 3 ligands and 252 TFs (39%) shared by any of 2 ligands (Figure 3A). Importantly, scalloped (sd), the downstream TFs of the Hippo/Yki signaling pathway was among the 640 TFs, (sup figure2), suggesting that sd and its co-fator Yki potentially regulate, at least in part, the expression of the secreted factors (Houtz *et al*. 2017).

**Figure 3:**
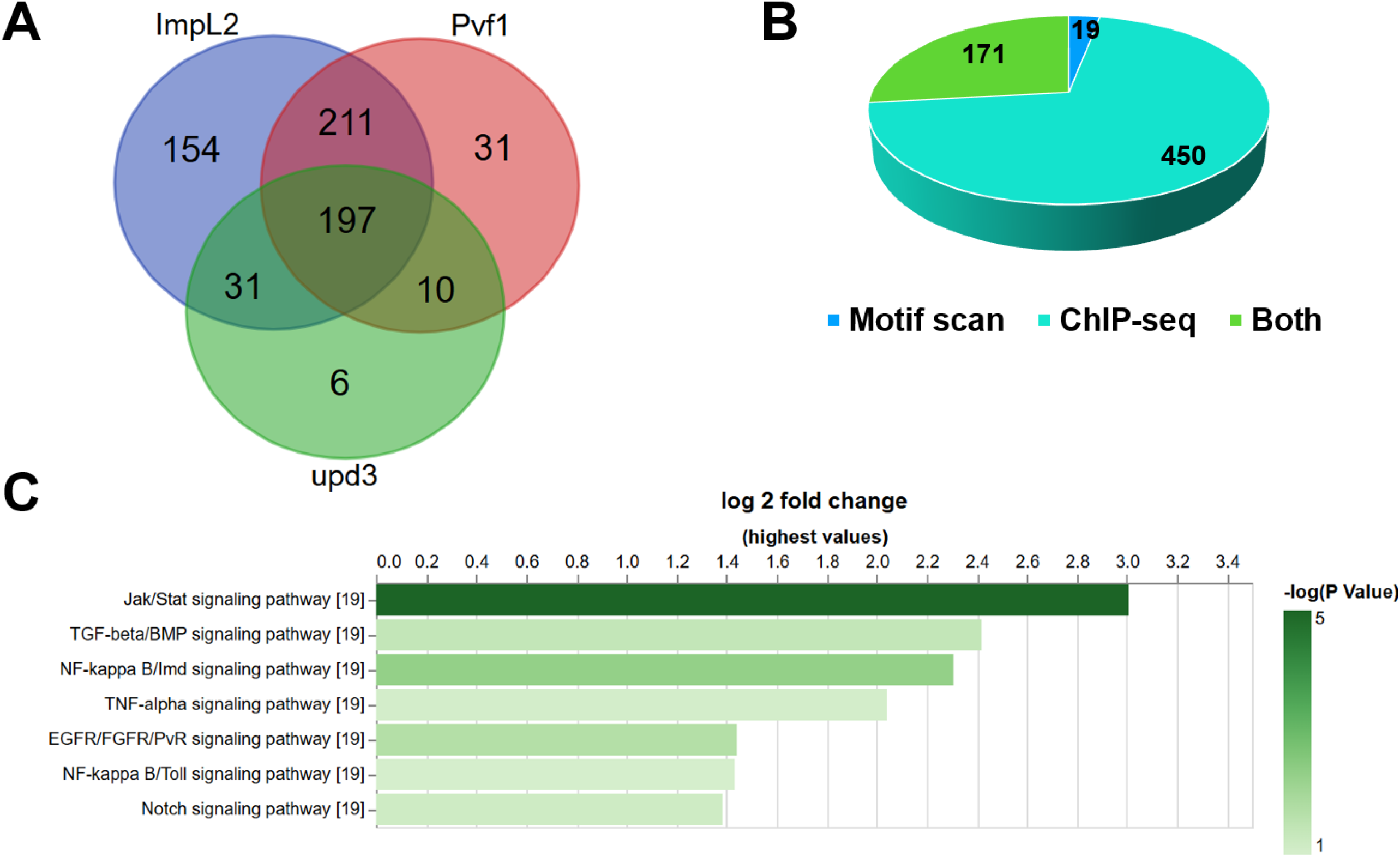
Identification of potential TFs that regulate the expression of cachexia ligands in Yki tumor using TF2TG. TFs for each ligand were searched at TF2TG with 5k search window and the combined. A. There is a significant overlap between the TFs for each ligand. B. Comparison of the TFs identified by motif scan and ChIP-seq data. C. Gene set enrichment analysis (GSEA) was performed on the combined TF list based on the core component analysis of major signaling pathway annotation using PANGEA.

The majority of the ChIP-seq datasets were obtained from embryos and larva (Kudron *et al*. 2018; Kudron *et al*. 2024). However, cachexia ligands were identified in the gut of adult flies bearing Yki tumors. Therefore, some of the TFs identified might not be expressed in the adult gut. To annotate the TFs accordingly, we retrieved the transcriptomic data obtained from the gut of 8 days old adult flies bearing Yki tumors (GSE113728) and found that 392 of the 640 TFs were expressed in that tissue, and therefore only focused the following analysis on these TFs. Next, we performed Gene Set Enrichment Analysis (GSEA) of these 392 TFs, based on protein complex annotation as well the core component annotation of major signaling pathways using PANGEA (Hu *et al*. 2023), to identify protein complexes and signaling pathway genes over-represented. The GSEA analysis revealed that Notch, Jak/Stat and EGFR/FGFR/PvR signaling pathways are activated in Yki gut tumors, which is consistent with the literature (Song *et al*. 2019; Ding *et al*. 2021; Pranoto *et al*. 2023). In addition, the Imd and TGF-beta pathways also scored both at the pathway and protein complex enrichment level (Figure 3C) (sup table 3). TFs of the TNF-alpha/NF-kappa B complexes that can potentially regulate the cachexia ligands are Rel (Relish), dl (dorsal) and Dif (Dorsal-related immunity factor), while the TFs of the SMAD/TGF-beta signaling complexes that can potentially regulate the ligands are Snoo (Sno oncogene) and Smox (Smad on X). In addition, we observed that the binding sites of different TFs are close to each other and overlap with annotated CRM regions. For example, the binding sites of sd, Rel, dl and Dif in the regulatory region of *upd3* overlap or are close to each other (sup figure2A). In addition, 5 of the 7 binding sites of these 4 TFs are within the same CRM (FBsf0000884519) as annotated by REDfly (sup figure2B), suggesting their potential co-regulatory functions. Interestingly, sd binding is observed in the regulatory sequences of all five TFs (sup table 2) suggesting an involvement of sd in regulating their expression. Signaling pathways are highly interconnected while the mutual regulation between Hippo-YAP pathway and innate immunity have been reported in mammals (Wang *et al*. 2020). However, in most biological systems, the mechanism of transcriptional regulation and how signaling pathways interact are still understudied. Insights gained from our TF2TG analysis identify potential TFs in Yki tumors that regulate the expression of cachexia-associated ligands as well as the potential cross-talk between the Hippo and other signaling pathways.

Using TF2TG of the cachexia ligands led to a number of predictions such as the involvement of IMD and TGF-beta signaling pathways regulating the expression of cachexia ligands. Indeed, the connection of IMD signaling with cachexia in Yki tumor model has been validated experimentally in a recent report (Singh *et al*. 2025), while TGF-beta pathway still needs to be tested. This use case demonstrates that TF2TG can assist researchers to identify TF candidates, while using transcriptomic data and protein complex annotation prioritize the TFs for formulating hypotheses about the transcriptional network as well as the relevant signaling pathways. In addition, the information of the potential binding sites at TF2TG can help scientists to design follow-up experiments and validate hypotheses.

### Concluding Remarks

There is a pressing need for tools that help researchers develop transcriptional networks and specific testable hypotheses regarding mechanisms of transcriptional regulation. Despite the availability of different types of relevant data (e.g. ChIP-seq, Hi-C, ChIA-PET, and ATAC-seq data as generated by 4DN and other efforts (Dekker et al. 2017), as well as the availability of data-informed consensus binding site motif prediction, there remains a need to integrate these data with the specific goals of developing transcriptional networks and using available data to predict what TFs might control transcription of a given gene of interest. TF2TG fills a currently unmet need by making it possible for researchers to input a gene and retrieve testable predictions, based on integrated sources, on the potential TFs that regulate that gene or to input a TF to identify the potential downstream target genes. Using TF2TG, we identified sd, a TF downstream of the Yki/Hippo signaling pathway, as a potential regulator of the Yki-induced cachectic secreted factors as well as the potential cross-talk between the Hippo pathway and other pathways such as TNF-alpha/NF-kappa B signaling and Smad/TGFbeta signaling in regulating the expression of the cachectic ligands Upd3, Pvf1 and ImpL2.

As more single-cell RNA seq (scRNA-seq) data sets become available and the cost of single-cell technology drops, scRNA-Seq will increasingly be used to study the state and dynamics of cells with high throughput and accuracy. The TF2TG resource will be well-positioned to help researchers make the most of scRNA-seq data sets to understand transcriptional regulation.

## Acknowledgements

We appreciate the feedback from the Perrimon lab for their valuable feedback. We extend our gratitude to the Harvard Medical School Research Computing and IT-Client Services teams for their consultation, web hosting, and support. This article is governed by HHMI’s Open Access to Publications policy. HHMI lab heads have previously granted a nonexclusive CC BY 4.0 license to the public and a sublicensable license to HHMI for their research articles. Accordingly, the author-accepted manuscript of this article will be made freely available under a CC BY 4.0 license immediately upon publication. This work was supported in part by a grant from the U.S. National Institutes of Health (NIH) National Institute of General Medical Sciences (P41 GM132087) that established our group as the Drosophila Research and Screening Center-Biomedical Technology Research Resource (DRSC-BTRR), as well as by grants from the NIH Office for Research Infrastructure Projects (R24 OD026435, R24 OD030002, R24 OD019847, R24 OD031952) to support resource development. In addition, M.L.B. is supported by R01 HG009723 and R56 HG009723 from NIH and N.P. is an investigator of Howard Hughes Medical Institute.

## Data Availability Statement

The resource is available to pubic and the URL is https://www.flyrnai.org/tools/tf2tg.

